# PPARγ activation by lipolysis-generated ligands is required for cAMP dependent *UCP1* induction in human thermogenic adipocytes

**DOI:** 10.1101/2024.08.10.607465

**Authors:** Anand Desai, Zinger Yang Loureiro, Tiffany DeSouza, Qin Yang, Javier Solivan-Rivera, Silvia Corvera

**Author notes:** **Corresponding author at:** Program in Molecular Medicine, University of Massachusetts Chan Medical School, Worcester, MA 01605, USA **Email:**. **Author Contributions:** A.D. and S.C. designed research; A.D. and Q.Y. and performed research; T.D., J.S.R. and Z.Y.L. contributed new reagents/analytical tools; A.D., Z.Y.L. and S.C. analyzed data; S.C. wrote the paper. **Competing Interest Statement:** The authors declare no competing interest.

## Abstract

**Objective:** The uncoupling protein 1 (UCP1) is induced in brown or “beige” adipocytes through catecholamine-induced cAMP signaling, which activates diverse transcription factors. UCP1 expression can also be enhanced by PPARγ agonists such as rosiglitazone (Rsg). However, it is unclear whether this upregulation results from de-novo differentiation of beige adipocytes from progenitor cells, or from the induction of UCP1 in pre-existing adipocytes. To explore this, we employed human adipocytes differentiated from progenitor cells and examined their acute response to Rsg, to the adenylate-cyclase activator forskolin (Fsk), or to both simultaneously.

**Methods:** Adipocytes generated from primary human progenitor cells were differentiated without exposure to PPARγ agonists, and treated for 3, 6 or 78 hours to Fsk, to Rsg, or to both simultaneously. Bulk RNASeq, RNAScope, RT-PCR, CRISPR-Cas9 mediated knockout, oxygen consumption and western blotting were used to assess cellular responses.

**Results:** *UCP1* mRNA expression was induced within 3 hours of exposure to either Rsg or Fsk, indicating that Rsg’s effect is independent on additional adipocyte differentiation. Although Rsg and Fsk induced distinct overall transcriptional responses, both induced genes associated with calcium metabolism, lipid droplet assembly, and mitochondrial remodeling, denoting core features of human adipocyte beiging. Unexpectedly, we found that Fsk-induced *UCP1* expression was reduced by approximately 80% following CRISPR-Cas9-mediated knockout of *PNPLA2*, the gene encoding the triglyceride lipase ATGL. As anticipated, ATGL knockout suppressed lipolysis; however, the associated suppression of UCP1 induction indicates that maximal cAMP-mediated *UCP1* induction requires products of ATGL-catalyzed lipolysis. Supporting this, we observed that the reduction in Fsk-stimulated UCP1 induction caused by ATGL knockout was reversed by Rsg, implying that the role of lipolysis in this process is to generate natural PPARγ agonists.

**Conclusion:** *UCP1* transcription is known to be stimulated by transcription factors activated downstream of cAMP-dependent protein kinases. Here we demonstrate that *UCP1* transcription can also be acutely induced through PPARγ-activation. Moreover, both pathways are activated in human adipocytes in response to cAMP, synergistically inducing UCP1 expression. The stimulation of PPARγ in response to cAMP occurs as a result of the production of natural PPARγ activating ligands through ATGL-mediated lipolysis.

**GRAPHICAL ABSTRACT:** 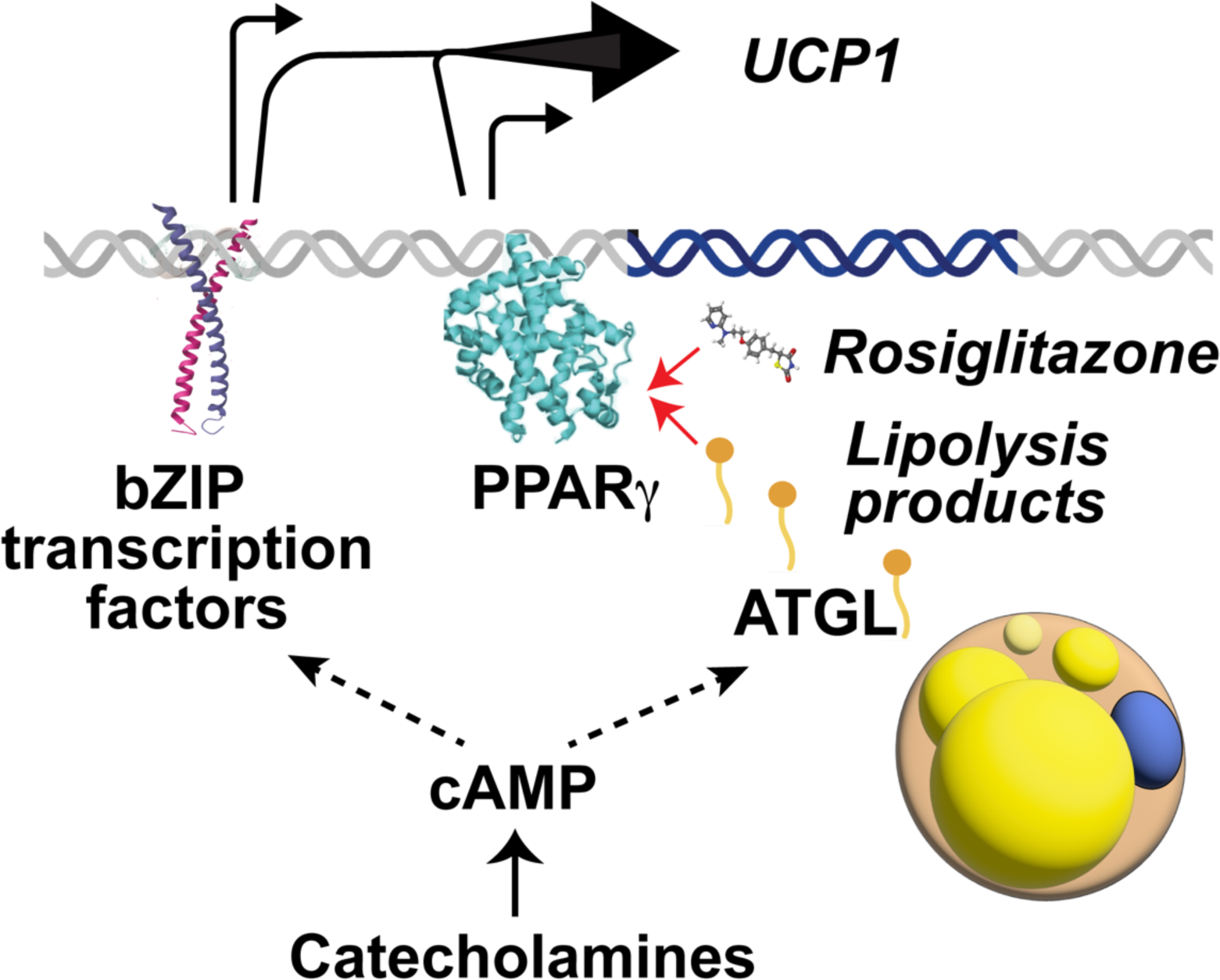

## INTRODUCTION

Adipose tissue plays key roles in systemic metabolism through multiple functions, including the storage of energy as triglycerides, the release of fatty acids for energy use during fasting, and non-shivering thermogenesis. The most extensively studied mechanism of adipocyte thermogenesis involves the expression of the uncoupling protein UCP1, which accelerates electron transport to produce heat [1]. UCP1-expressing adipocytes comprise most of the brown adipose depot located within the interscapular region of rodents and are also found distributed among white adipocytes within inguinal mouse fat, where they are referred to as brite/beige adipocytes [2]. In humans, UCP1-containing adipocytes are interspersed within supraclavicular, paravertebral and perirenal adipose tissue depots, resembling the distribution of beige adipocytes in mice [3]. Understanding the mechanisms underlying the development and maintenance of UCP1-expressing adipocytes in humans has gained interest due to the strong inverse correlation between thermogenic adipose tissue abundance and metabolic disease risk [4]. Additionally, the implantation of human UCP-1-expressing adipocytes into mice can mitigate obesity-associated metabolic disfunction [5–7], supporting the direct impact of these cells on metabolic health.

Most research on the mechanism regulating thermogenesis have been conducted in mice, where *Ucp1* transcription is stimulated by catecholamines released upon cold exposure [8]. Catecholamines trigger cAMP production, which activates PKA-mediated phosphorylation of multiple targets, resulting in activation of bZIP transcription factors that can induce UCP1 [9–11]. In addition, cAMP induces lipolysis, which produces fatty acids required for activation of UCP1 uncoupling in mitochondria [12]

In addition to catecholamines, PPARγ agonists such as rosiglitazone and pioglitazone increase expression of UCP1 in adipose tissues of mice [13–15], and in primary adipocyte cultures [16–18]. However, it has been difficult to distinguish whether PPARγ activation increases UCP1 levels by increasing the number of beige adipocytes differentiating from progenitor cells, or whether it acts on fully differentiated adipocytes to induce UCP1 expression. Indeed, PPARγ activation could bias adipocyte differentiation by antagonizing Wnt signaling, leading to unbalanced development of distinct adipocyte subtypes [19, 20]. Because most established protocols for primary mouse or human adipocyte differentiation require the use of PPARγ agonists, the acute effects of these agonists on differentiated primary adipocytes have been difficult to assess.

In prior work, we have found conditions to generate human adipocytes from multipotent mesenchymal progenitor cells without the need for PPARγ agonists during the differentiation process[6, 20–22]. We derive progenitor cells from human adipose tissue fragments cultured in MatriGel with pro-angiogenic medium. Progenitor cells derived by this method can differentiate into multiple adipocyte subtypes, including beige adipocytes [21]. Here, we have leveraged this approach to study the mechanisms by which cAMP and PPARγ activation induce *UCP1* in human adipocytes. We find that each pathway can independently induce *UCP1* transcription, and that they act cooperatively for maximal induction. Importantly, we find that cAMP can activate both pathways, activating PPARγ through ATGL-dependent activation of lipolysis.

## RESULTS

We have previously reported that human multipotent mesenchymal progenitor cells can be obtained in large numbers by culturing adipose tissue fragments under pro-angiogenic conditions [23]. Progenitor cells derived by this method differentiate efficiently using a minimal cocktail of methyl isobutyl-xanthine, dexamethasone and insulin (MDI, Fig. 1), evidenced by the appearance of lipid droplets in most cells (Fig.1 a,b), and by induction of canonical adipocyte marker genes (Fig 1c). To determine whether adipocytes differentiated in this way would respond to acutely rosiglitazone (Rsg), we measured *UCP1* mRNA. *UCP1* was induced by Rsg as early as 3h (Fig.1d), an effect that cannot be accounted for by an increased number of differentiated cells, as other canonical adipogenic genes were not induced (Fig. 1e). As expected, cells also responded robustly to Fsk with *UCP1* induction within 3h of exposure (Fig. 1f).

**Figure 1.**
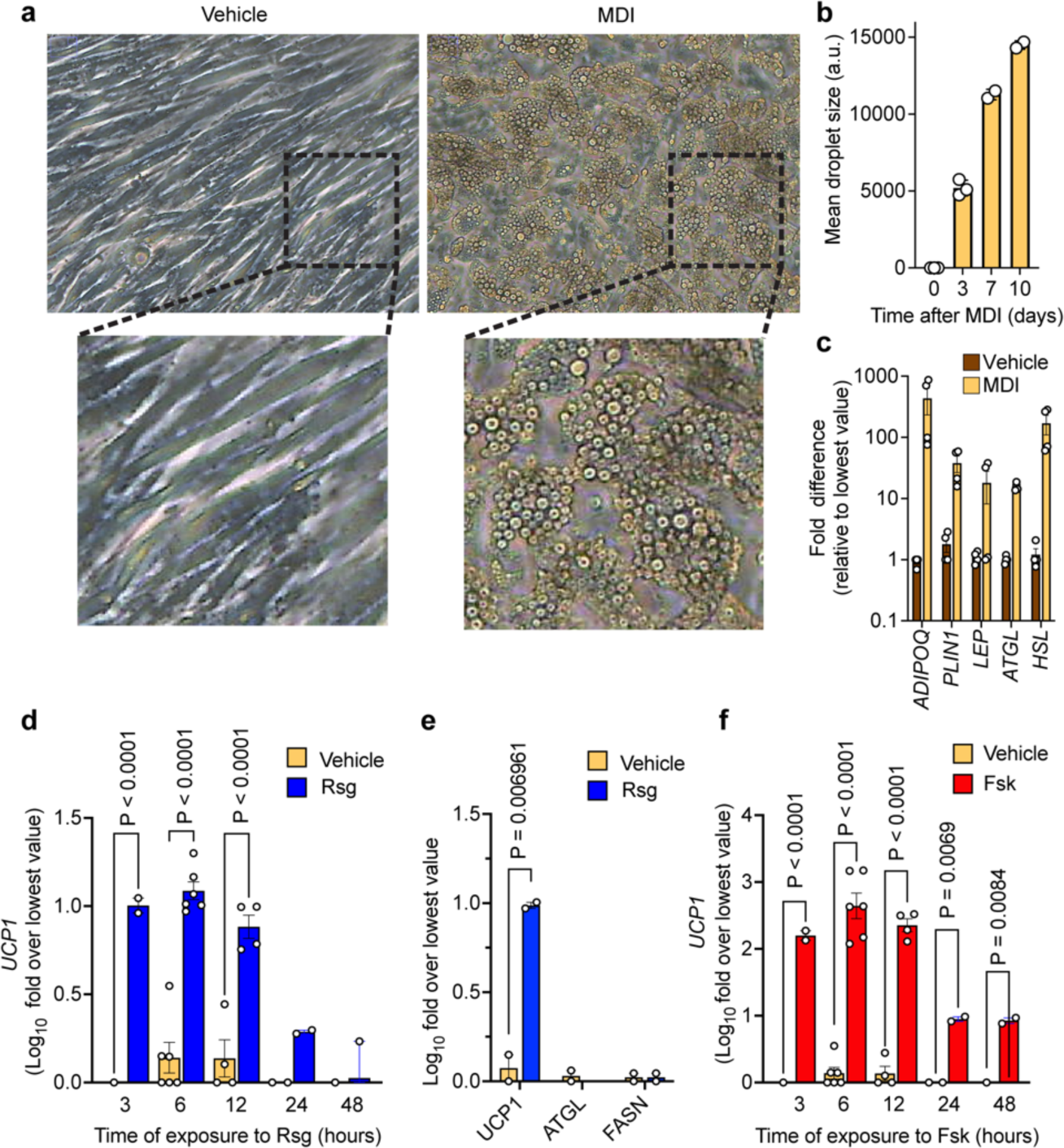
Features of human adipocytes differentiated from progenitor cells without added PPARγ ligands. Progenitor cells were exposed to vehicle or differentiation cocktail (MDI) and analyzed after 10 days (**a,c-f**), or at the times indicated (**b**). **a**) Phase images of cells. **b**) Mean droplet size. **c**) mRNA levels of genes indicated in the x-axis. **d)** mRNA levels of UCP1 in response to 1μM Rsg for the times indicated. **e)** mRNA levels of genes indicted in the x-axis after 6h of exposure to 1μM Rsg. **f)** mRNA levels of UCP1 in response 10μM Fsk for the times indicated. Plotted are the means and SEM of independent culture wells, derived from a minimum of 2 subjects, with each value shown. Significance of the differences between conditions was calculated using ANOVA after multiple comparison testing, and exact P-values are shown.

In addition to *UCP1* expression, the establishment of a thermogenic phenotype requires an increase in mitochondrial density and formation of multilocular lipid droplets. To determine whether the induction of *UCP1* by Rsg or Fsk entailed induction of a shared thermogenic gene expression program, we conducted bulk RNA Seq. After 6 hours of stimulation by Rsg, 29 genes were upregulated, and 4 genes downregulated with a P-adj value <0.05 and fold effect > 2 (Fig. 2a, and ntary Table 1). The top 20 biological processes significantly enriched by upregulated genes included peptide and metabolite responsiveness, adaptive thermogenesis, response to lipids, and brown fat cell differentiation (Fig. 2b). Fsk produced a much more robust response, with 657 genes upregulated and 501 downregulated after 6h of exposure, with a P-adj value <0.001 and 2x fold effect (Fig. 2c, Supplementary Table 1). The top 20 processes enriched by Fsk-upregulated genes included hormonal responsiveness, fat cell differentiation and response to lipid (Fig. 2d).

**Figure 2.**
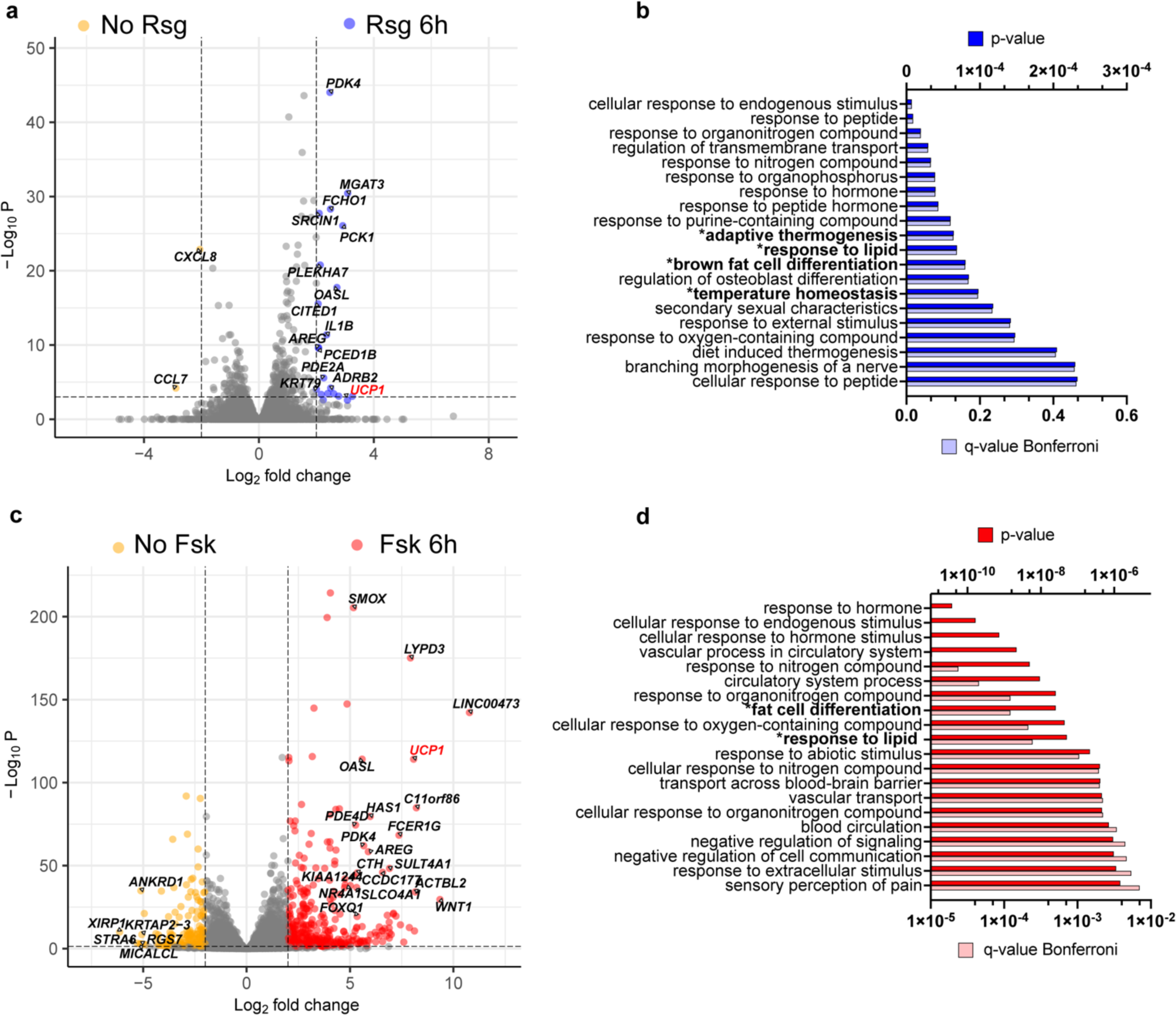
Transcriptional responses and enrichment analysis of human adipocytes exposed acutely to Rsg or Fsk. **a,c**. Volcano plots of genes differentially expressed between adipocytes treated for 6h with either 1μM Rsg (a) or 10μM Fsk (c). **b,d.** GO: Biological Processes enriched by genes upregulated by Rsg (b) or Fsk (d), using ToppGene [49].

A larger number of genes were affected after longer exposure (78 hours) to Rsg, with 349 genes upregulated, and 62 downregulated with a P adj value < 0.001 and >2-fold effect (Fig. 3a, Supplementary Table 1). The top 20 pathways enriched by upregulated genes were primarily associated with lipid metabolism (Fig. 3b). Interestingly, the number of gene affected after longer exposure to Fsk was similar to that seen after only 6h, with 625 genes were upregulated, and 618 downregulated (Fig 3c, Supplementary Table 1. The top 20 pathways enriched by Fsk-upregulated genes were primarily associated with lipid metabolism (Fig 3d), similar to that seen in response to Rsg. These results indicate that, while transcriptional responses to Rsg and Fsk differ markedly in kinetics and number of affected genes, both acutely induce *UCP1* together with genes associated with lipid metabolism.

**Figure 3.**
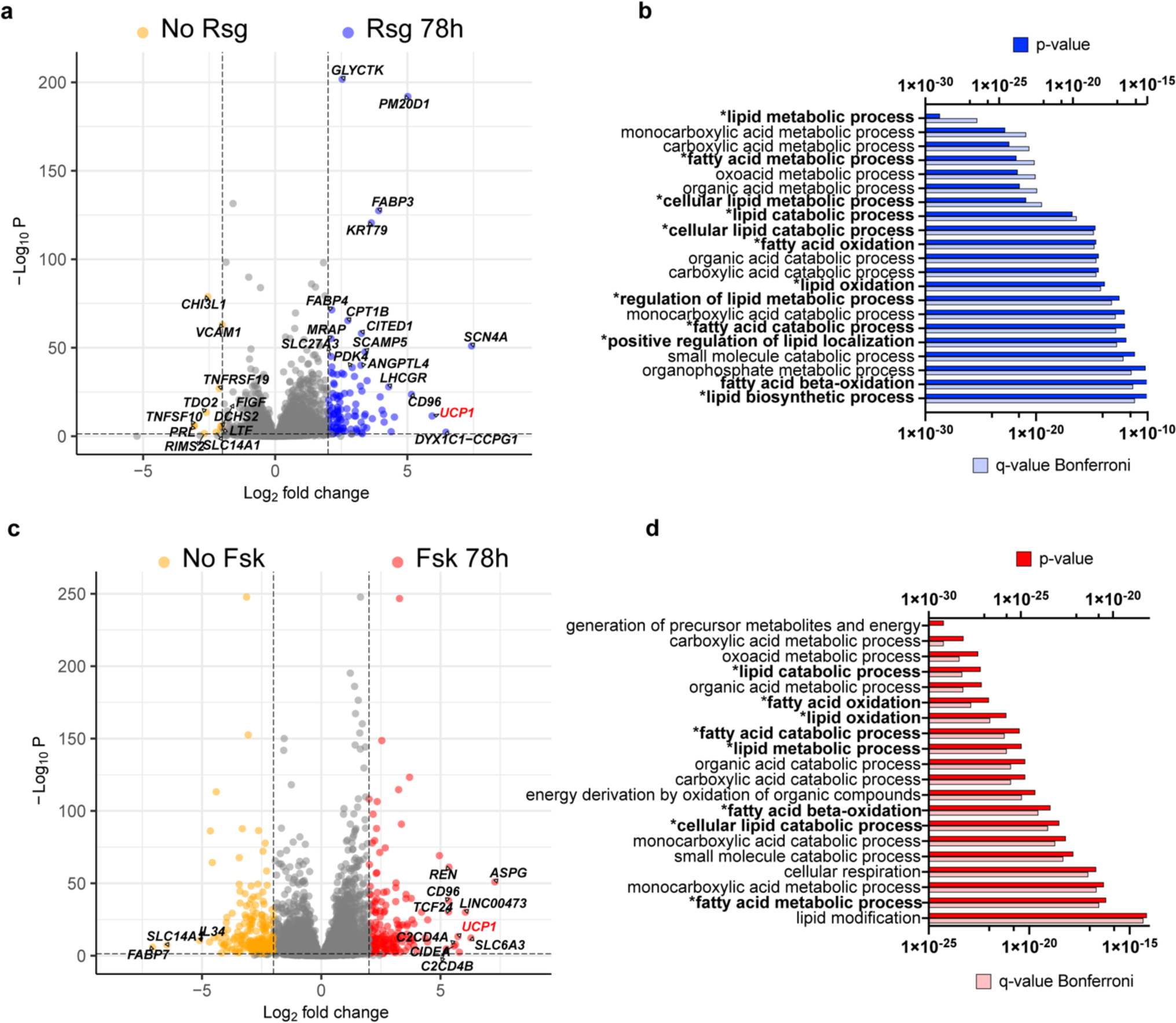
Transcriptional responses and enrichment analysis of human adipocytes after prolonged exposure to Rsg or Fsk. **a,c**. Volcano plots of genes differentially expressed between adipocytes treated for 78 hours with either 1μM Rsg (**a**) or 10μM Fsk (**c**). Doted lines are set at P adj value <0.001 (-Log10) and fold effect > 2 or < 2. **b,d**. GO: Biological Processes enriched by genes upregulated by Rsg (b) or Fsk (d), using ToppGene [49].

To identify genes that might represent the core components of the thermogenic phenotype, we filtered the data to identify those upregulated by both Rsg and Fsk (Supplementary table 1). After 6h of stimulation, 13 genes were upregulated by both Fsk and Rsg by a minimum of 2-fold, with a P adjusted value < 0.05 (Fig. 4a). Pathways enriched by these genes included calcium transport, thermogenesis, and G-protein coupled receptor signaling (Fig. 4b). After 78h of stimulation, 144 genes were upregulated by both Fsk and Rsg (> 2-fold, P adjusted value < 0.001) (Fig. 4c). These genes were associated with fatty acid oxidation, triglyceride biosynthesis, hormonal responsiveness, and thermogenesis (Fig. 4d, Supplementary table 1). Moreover, these genes were associated with specific cellular compartments, predominantly lipid droplets and the mitochondrial matrix (Fig. 4e). These results suggest that hormonal signaling pathways and calcium transport are necessary early upon induction of the thermogenic phenotype, followed by major adaptations in lipid dynamics and organelle architecture.

**Figure 4.**
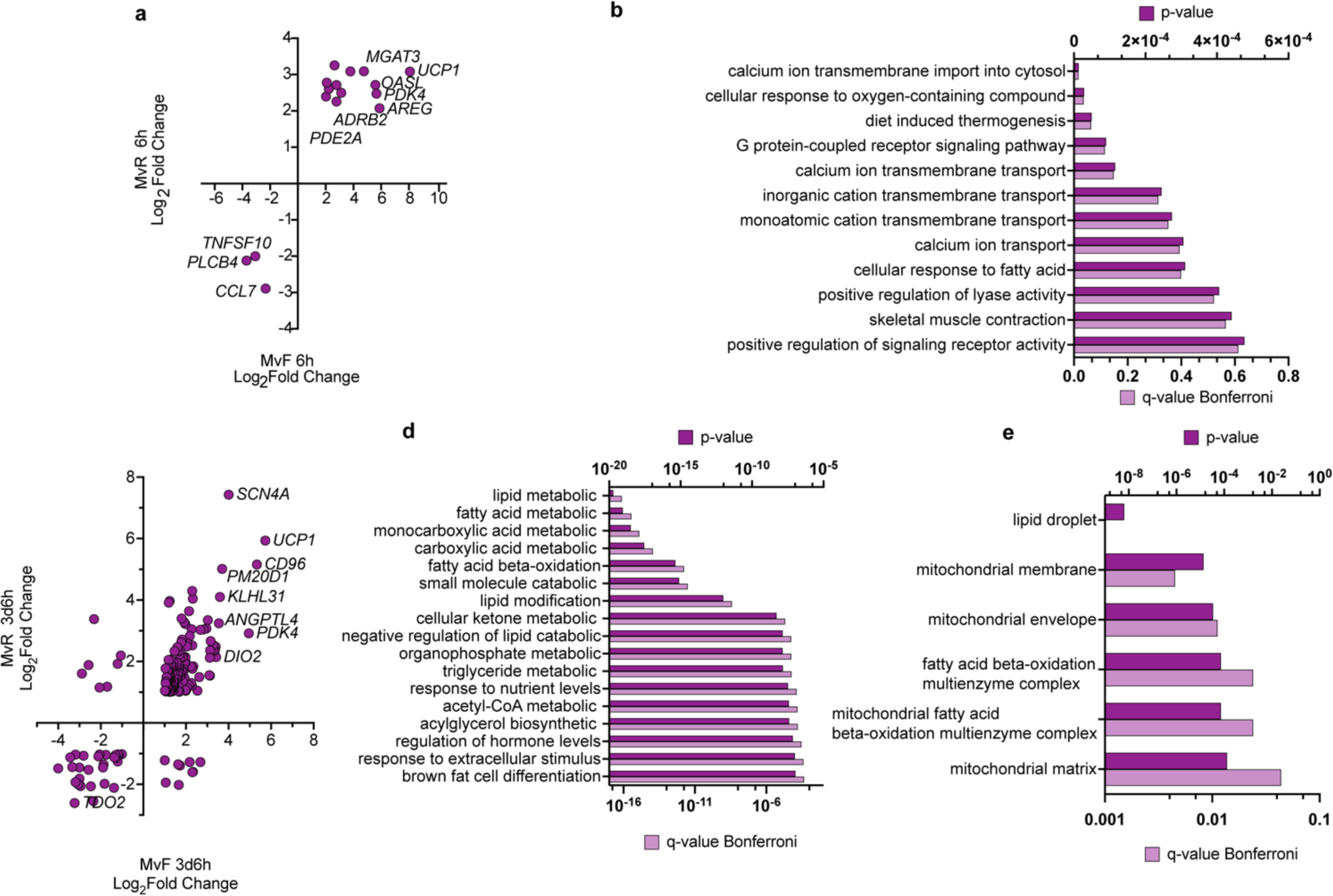
Common transcriptional responses between Rsg and Fsk. **a, c.** Upper right and lower left quadrants indicating genes similarly regulated by both Fsk (x-axis) and Rsg (y-axis) after 6 (a) or 78 (c) hours of exposure. **b,d**. Biological processes enriched by genes affected by both Fsk and Rsg after 6 (b) or 78 (d) hours of exposure. **e.** Cellular components enriched by genes affected by both Fsk and Rsg after 78 hours of exposure.

Human progenitor cells can differentiate into various types of adipocytes [21, 24], raising the possibility that Rsg and Fsk might act on distinct adipocyte subtypes to induce *UCP1*. To explore this possibility, we used RNA Scope to detect *UCP1* mRNA in individual cells (Fig. 5a). A clear signal was detected in numerous cells in response to either Rsg or Fsk. However, when Rsg and Fsk were added together, we detected a much more intense signal in many cells, indicating that these ligands stimulate *UCP1* expression in the same adipocyte. Further quantification confirmed a large cooperative effect of Rsg and Fsk on *UCP1* induction (Fig. 5b), which was also evident by bulk analysis using RT-PCR (Fig. 5c)

**Figure 5.**
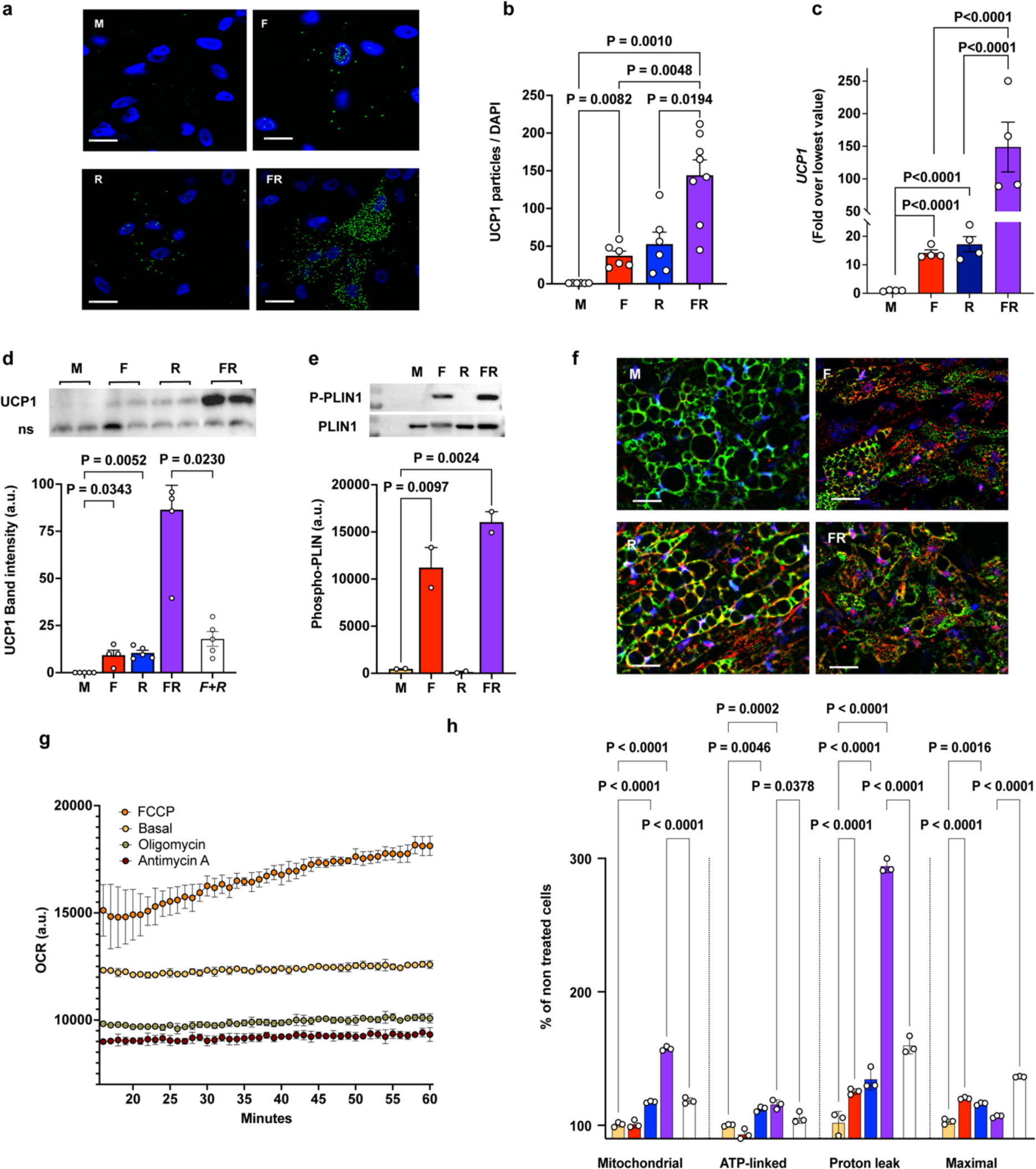
Synergistic induction of UCP1 by Rsg and Fsk. Adipocytes differentiated for 10 days were treated with vehicle (M, yellow bars), 10μM Fsk (F, red bars), 1μM Rsg (R, blue bars), or Fsk and Rsg simultaneously (FR, purple bars) for 78h. Clear bars represent the arithmetic sum of the effects of F and R applied individually, enabling evaluation of any synergistic effect occurring when F and R are applied simultaneously. **a.** Visualization of *UCP1* mRNA in individual cells using RNAScope (green) and Hoescht nuclear staining (blue). **b.** Quantification of number of particles per cell in 6-8 fields, expressed as intensity of *UCP1* mRNA staining divided by intensity of Hoescht staining per field. **c.** *UCP1* mRNA quantified by RT-PCR from four independent experiments. **d.** Example of western blotting with an antibody to UCP1, and quantification of band intensities from 4 independent experiments. **e.** Examples of western blotting with antibodies to phospho-PLIN1, or total PLIN1as indicated, and quantification of phospho-PLIN1 from two independent experiments. **f.** Examples of immunofluorescence staining with antibodies to PLIN1(green), UCP1 (red) and Hoescht nuclear staining (blue). **g.** Oxygen consumption rate over time of cells (M) treated with the indicated reagents. **h.** Oxygen consumption parameters, where Mitochondrial=basal-antimycin, ATP-linked=basal-oligomycin, Uncoupled=oligomycin-antimycin and Maximal=FCCP-antimycin. In each experiment, the mean of last 5 measurements of the OCR time course for each treatment conditions were used to calculate parameters, and plotted are n=3 independent experiments. For b-e,h plotted are means and SEM of each independent experiment, and statistical significance calculated using ANOVA corrected for multiple comparisons with exact P-values shown.

We then addressed whether the synergistic induction of *UCP1* mRNA would translate into UCP1 protein expression and function, and its potential mechanism. Western blotting revealed that UCP1 protein levels were increased by either Fsk or Rsg, and synergistically by their combination (Fig. 5d). The synergistic effects of Fsk and Rsg effect could not be explained by alterations in cAMP signaling, as Rsg alone had no effect, nor did it significantly enhance, the effect of Fsk on PLIN1 phosphorylation (Figure 5e). The absence of an effect of Rsg on cAMP-mediated signaling was also evidenced by analysis of lipid droplet morphology, as droplet size decreases markedly in response to Fsk, but not in response to Rsg, despite both inducing UCP1 protein (Fig. 5f).

To assess whether UCP1 was functional, we measured oxygen consumption under basal conditions and after exposure to antimycin, oligomycin, or FCCP. We used a method that enables monitoring oxygen consumption by cells differentiated in 96-well multiwell plates exposed individually to each agent, minimizing mechanical disruption of attached adipocytes (Fig. 5g). We then calculated mitochondrial (basal-antimycin), ATP linked (basal-oligomycin), uncoupled (oligomycin-antimycin) and maximal (FCCP-antimycin) oxygen consumption from equilibrium values (mean of last 5 measurements) in all treatment conditions (Fig. 5h). We find that exposure of adipocytes to Rsg increased mitochondrial, ATP-linked, maximal and uncoupled oxygen consumption compared to untreated cells, consistent with a larger overall mitochondrial capacity and UCP1-mediated uncoupling. Exposure of adipocytes to Fsk increased maximal and uncoupled oxygen consumption. Exposure to both Fsk and Rsg resulted in a large synergistic increase in uncoupled respiration, consistent with the observed increase in UCP1 protein. These results indicate that the synergistic induction of *UCP1* mRNA by Rsg and Fsk leads to accumulation of functional UCP1 protein and mitochondrial uncoupling.

To explore the mechanism underlying the synergistic effects of Rsg and Fsk on *UCP1* mRNA and protein levels, we first asked whether other genes responsive to Fsk would also be synergistically induced by Fsk and Rsg. *LINC00473*, a highly responsive gene specific to human beige adipocytes [25], was induced by Fsk but not Rsg, and Rsg suppressed the effect of Fsk (Fig. 6a). These results indicate that the synergistic effect of Rsg and Fsk on UCP1 expression is not shared by all Fsk-responsive genes. To identify genes that are similarly regulated, we performed RNA-Seq on cells stimulated with either Rsg (R), Fsk (F) or Rsg and Fsk together (RF), and filtered the data to find genes stimulated synergistically, i.e. where expression in RF would be larger than the arithmetic sum of R+F. Strikingly few genes were synergistically induced; indeed, UCP1 was conspicuous with a RF/R+F of 4.51, while the mean of all other synergistically upregulated genes was 1.47 ± 0.6 (Fig. 6b).

**Figure 6.**
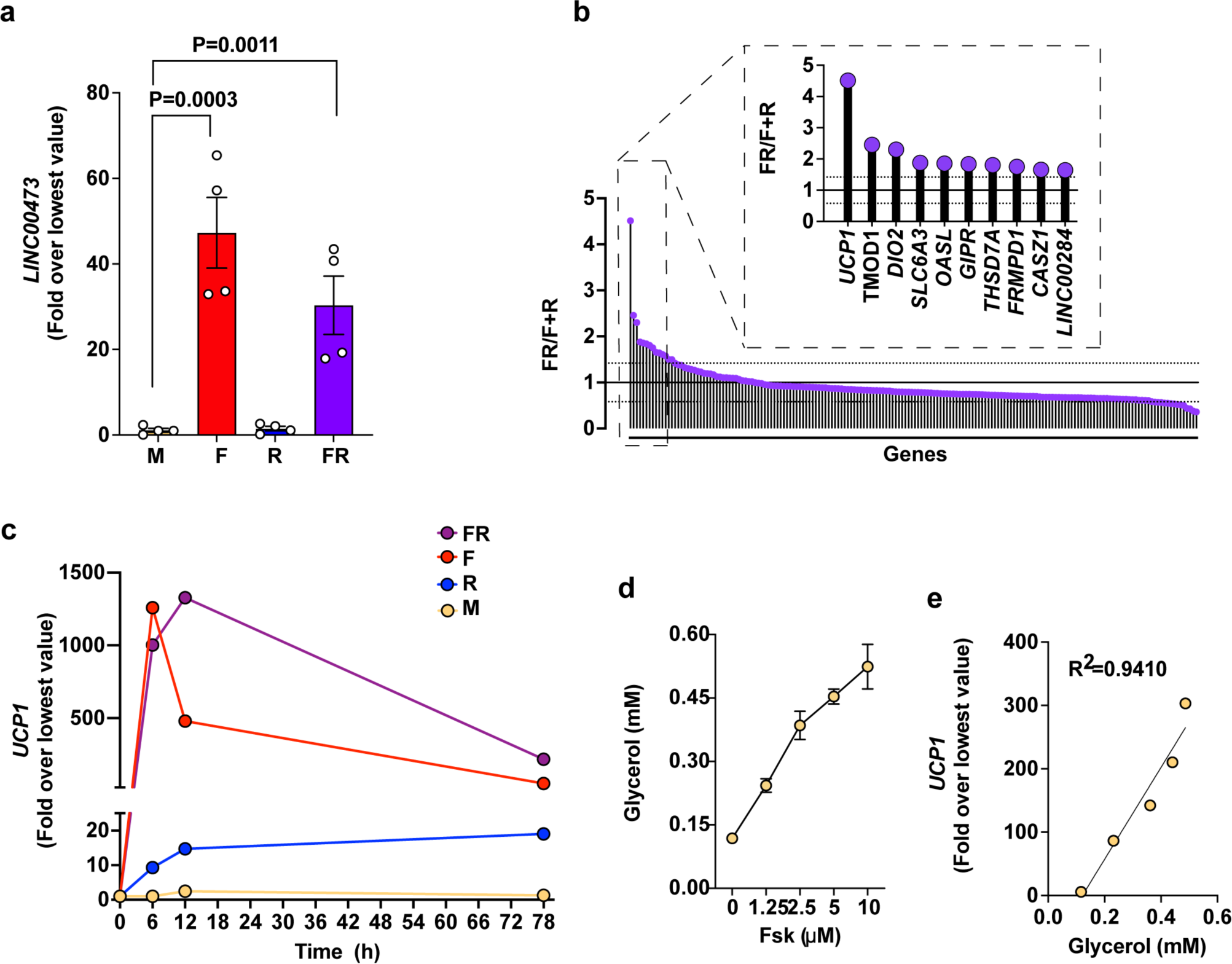
Only few genes are synergistically induced by Rsg and Fsk. Cells were differentiated for 10 days and treated with vehicle (M, yellow bars and symbols), Fsk (F, red bars and symbols), Rsg (R, blue bars and symbols), or Fsk and Rsg simultaneously (FR, purple bars and symbols) **a.** Expression of *LINC00473* after 78 h of treatment. Bars represent means and SEM of experiments with cells from two subjects, performed in duplicate. Statistical significance was calculated using ANOVA, and exact P-values are shown. **b**. TPM values from RNASeq datasets from cells treated for 78h with F, R or FR were used to calculate the ratios of the effect of FR (10μM Fsk and 1μM Rsg applied simultaneously) over the arithmetic sum of the effects of Fsk and Rsg applied individually. A value of 1 represents no synergy. Top 10 values are highlighted. **c.** Time course of *UCP1* mRNA expression in response to the indicated treatments. Plot represents one experiment, where symbols are the mean of replicate samples, and similar results were observed in two additional experiments **d.** Glycerol concentration in the medium of cells treated for 6h with the indicated concentrations of Fsk. **e.** Relationship between glycerol concentration in the medium and UCP1 mRNA expression in cells treated for 6h with 0-10μM Fsk for 6h.

To better understand the features of the effects of Rsg and Fsk, we analyzed the temporal dynamics of *UCP1* induction (Fig. 6c). Induction of *UCP1* by Fsk was rapid, reaching maximal levels at 6h, and was followed by a rapid decline. In contrast, induction by Rsg was rapid, apparent within 6h, but was sustained. The synergistic effect of Fsk and Rsg was most apparent at 12h suggesting that Rsg might prolong the effects of Fsk signaling. In addition to its effects to activate transcription factors, cAMP signaling strongly stimulates lipolysis through PKA-mediated phosphorylation of PLIN1 and subsequent activation of ATGL [26]. Markussen et al [27] have shown that lipolysis per-se activates many the transcriptional responses attributed to cAMP signaling. To explore the potential role of lipolysis, we analyzed the association between glycerol release and UCP1 induction. We observed a dose-dependent stimulation of lipolysis within 6h of addition of Fsk (Fig. 6d), which was closely correlated with the extent of *UCP1* induction (Fig. 6e).

To directly determine whether lipolysis influenced Fsk-stimulated UCP1 induction, we employed CRISPR-Cas9 to delete *PNPLA2*, which encodes ATGL. We used a previously developed method [5] to electroporate ribonucleoprotein particles containing a selection of sgRNAs directed to the *PNPLA2* locus into progenitor cells. Cells were then differentiated for 10 days, and ATGL protein quantified by western blotting. We selected sgRNAs that produced undetectable levels of ATGL (Fig. 7a). without affecting HSL (Fig. 7b). The functional effects of ATGL knockout were apparent as early as 4 days after induction of differentiation, with the size of emerging lipid droplets being significantly larger compared to cells in which cells were electroporated with non-targeting sgRNAs (Fig. 7c), consistent with an inhibition of basal lipolysis.

**Figure 7.**
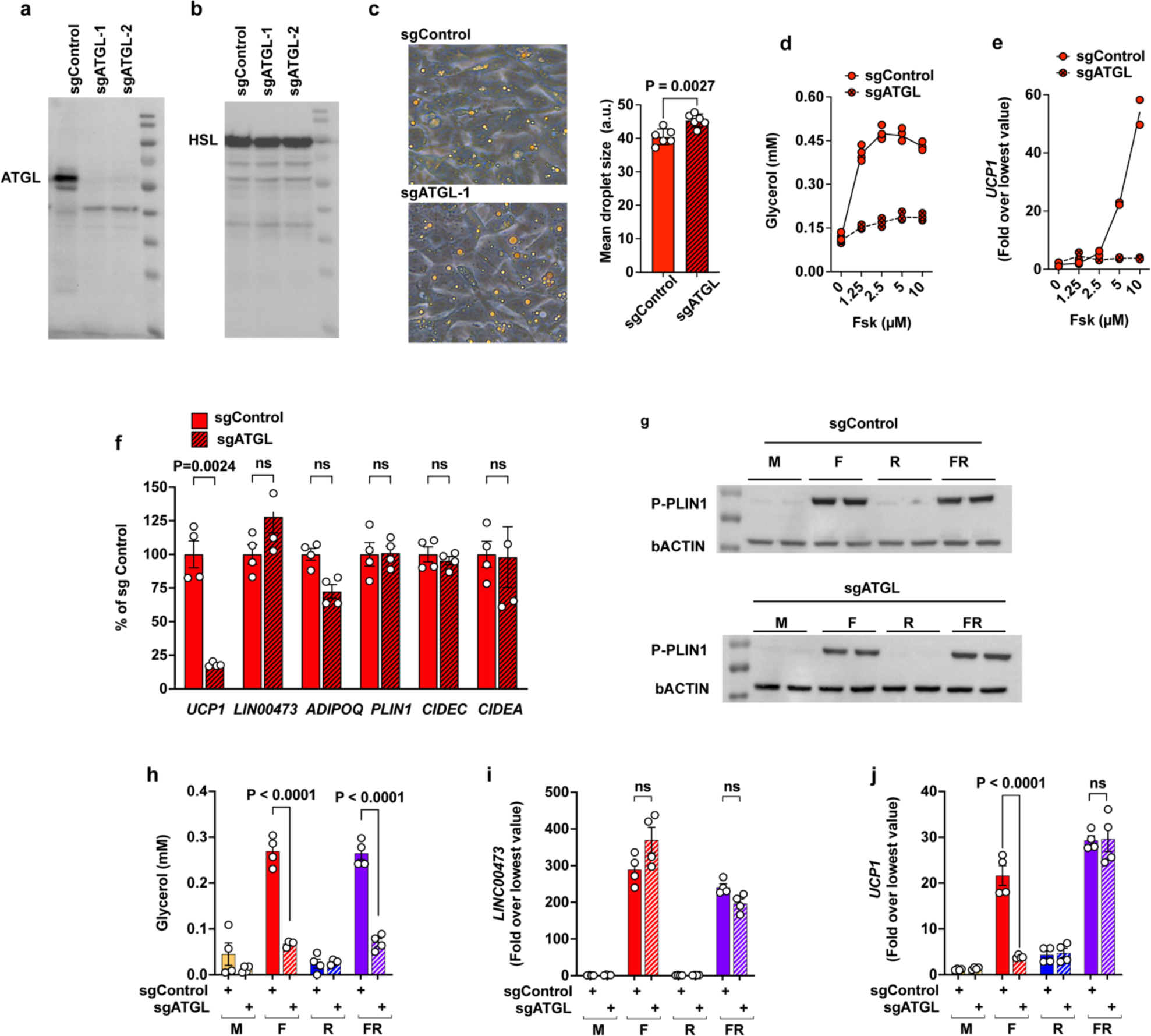
Induction of UCP1 by Fsk requires products of lipolysis. **a,b** Western blots using antibodies to ATGL (a), or HSL (b) of samples from cells electroporated with non-targeting (sgControl) or PNPLA2-targeting (sgATGL) RNA guides prior to differentiation for 10 days. **c.** Phase images of cells electroporated with non-targeting (sgControl) or PNPLA2-targeting (sgATGL) RNA guides after 4 days of differentiation, and quantification of lipid droplet size. **d-e.** Glycerol concentration in the medium (d), and UCP1 mRNA expression (e) of control or ATGL knockout cells differentiated for 10 days and treated for 6h with the indicated concentrations of Fsk. **f.** Expression of genes indicated in the x-axis, in control (sgControl, red bars), or ATGL knockout (sgATGL, red stripped bars) cells after 10 days of differentiation and 6h stimulation with 10 μM Fsk. Bars represent means and SEM of n=4 independent experiments. Values are calculated as a percent of expression in control cells. Statistical significance of differences was calculated using ANOVA and exact P values shown. g. Western blots using antibodies to phospho-PLIN1 and beta-actin as loading control. **h-j**. Glycerol concentration in the media (h), mRNA levels of LINC00473 (i), or mRNA levels of UCP1 (j), in control (sg Control, solid colors) or ATGL knockout (sgATGL, striped colors) cells, treated with vehicle (M, yellow bars), Fsk (F, red bars), Rsg (R, blue bars), or Fsk plus Rsg simultaneously (FR, purple bars). Bars represent means and SEM of n=4 independent experiments. Statistical significance of differences was calculated using ANOVA and exact P-values shown.

After 10 days of differentiation, glycerol release in response to Fsk was inhibited by ∼80% in ATGL knockout adipocytes (Fig. 7d). Strikingly, the effect of Fsk to induce *UCP1* was also strongly inhibited (Fig. 7e). In contrast, the effects of Fsk on other cAMP-sensitive genes including *LINC00473*, and other adipogenesis associated genes was not affected (Fig. 7f). Additionally, stimulation of PLIN1 phosphorylation by Fsk was similar between control and ATGL knockout cells (Fig. 7g). These results indicate that knockout of ATGL does not impair cAMP-dependent signaling, while, as expected, impairing lipolysis. Notably, these results reveal that induction of *UCP1* in response to Fsk in human beige adipocytes is highly dependent on ATGL-catalyzed lipolysis.

We then examined the effect of ATGL knockout on the actions of Rsg, and on the synergistic effects of Rsg and Fsk. As expected, Rsg alone did not stimulate lipolysis, and did not affect Fsk-stimulated lipolysis, in either ATGL knockout or control cells (Fig. 7h). Similarly, the stimulatory effect of Fsk on *LINC00473* was not affected by Rsg in either ATGL knockout or control cells (Fig. 7i). However, Rsg completely rescued the inhibitory effects of ATGL knockout on Fsk-stimulated *UCP1* induction (Fig. 7j). The finding that the inhibitory effect of ATGL knockout is reversed by Rsg strongly suggests that lipolysis produces natural activating ligands for PPARγ, and this activation cooperates with other branches of cAMP signaling to fully induce *UCP1* in human beige adipocytes.

## DISCUSSION

Catecholamine mediated, cAMP-dependent induction of UCP1 has been extensively studied as a major mechanism of non-shivering thermogenesis. However, it has also been recognized that browning can be achieved by non-adrenergic mechanisms which are likely to be functionally important, as in early development brown adipose tissue is recruited prior to adrenergic signaling [13, 17, 28]. Non-adrenergic mechanisms of browning have been traced to activation of PPARγ [13]. However, understanding the mechanism of PPARγ action is complicated by the broad, pro-adipogenic effect of this nuclear receptor, which is difficult to deconvolve from effects unrelated to adipogenesis per se. Here we report that PPARγ stimulation can acutely induce UCP1 transcription independently of new adipocyte recruitment.

The acute effects of Fsk and Rsg to induce *UCP1* are accompanied by broader transcriptional responses. We reasoned that the common elements between the Fsk and Rsg responses might represent the minimum complement of genes driving the cellular adaptations required for a sustained thermogenic response. Interestingly, at early time points, genes commonly induced by Fsk and Rsg enrich pathways associated with calcium signaling and hormonal responsiveness, and at later time points they enrich pathways associated with lipid metabolism and organelle architecture. These common adaptations are potentially also required to accommodate UCP1-independent thermogenic mechanisms that occur in brown adipocytes [29, 30], which are likely to require features of mitochondria and lipid droplets required to support fuel oxidation.

Our finding of a synergistic induction of *UCP1* when cells are simultaneously exposed to Fsk and Rsg is consistent with prior results in both cells and mice [14, 31–34], but mechanisms have remained unclear. The observation that this synergy is most pronounced during the decline phase of Fsk stimulation suggested that it may be attributable to prolongation of an acute effect of Fsk. Fsk acts through elevation of cAMP, which induces *UCP1* through activation of diverse transcription factors [35]. However, elevation of cAMP in adipocytes also induces hydrolysis of lipid droplet triglycerides through a process mediated sequentially by the three lipolytic enzymes– adipose triglyceride lipase (ATGL, *PNPLA2*) [36], hormone-sensitive lipase (HSL, *LIPE*) and monoacylglycerol lipase (MGL, *MGLL*) [37]. In response to cAMP signaling, PLIN1 becomes phosphorylated by protein kinase A (PKA), releasing CGI-58 which can then activate ATGL to initiate lipolysis [38, 39]. Our data showing that knockout of ATGL greatly reduces *UCP1* induction demonstrates that lipolysis is essential for the maximal response to cAMP.

PPARγ is activated by synthetic and natural ligands [40], which can co-bind to modulate its activity [41]. Although various natural products, including fatty acids and prostaglandins have been shown to bind and activate PPARγ in-vitro [42], determining their physiological relevance is challenging [43]. The discovery that a product of lipolysis can phenocopy Rsg, a synthetic activating ligand, suggests that natural PPARγ ligands are produced in human adipocytes through ATGL-mediated hydrolysis, paving the way for their identification in future studies. The synergistic effect on UCP1 induction that occurs in response to cAMP signaling and strong PPARγ agonists such as Rsg can potentially be explained by prolongation of the effect of natural PPARγ ligands produced through ATGL-catalyzed hydrolysis.

The large synergistic effect of cAMP-activated transcription factors and lipolysis is also evidenced in studies in mouse adipocytes. For example, in murine brown adipocyte cell lines, activation of lipolysis alone, achieved by direct modulation of CGI-58, had a much smaller effect on UCP1 transcription compared to that of isoproterenol, which stimulates cAMP signaling [27]. Moreover, inhibition of bZIP transcription factor activation without inhibition of lipolysis greatly reduced UCP1 induction in response to adrenergic stimulation [44, 45].Together these results indicate that activation of cAMP responsive transcription factors synergizes with PPARγ to induce UCP1 in multiple experimental models. While complex, these mechanisms could potentially be leveraged to enhance the functional capacity of thermogenic adipocytes in vivo [31].

## METHODS

### Cells

Progenitor cells were obtained from human abdominal subcutaneous adipose tissue explants and expanded according to previously published methods. Briefly, adipose tissue explants from patients undergoing panniculectomy procedures at University of Massachusetts Medical Center with the approval of University of Massachusetts Institutional Review Board (#14734_13) were embedded in Matrigel (Corning 356231; 200 explants with sizes of 1cm^3^ per 10 cm dish) and cultured in angiogenic conditions for 14 days. Progenitor cells sprouting from the explants were obtained using Dispase (Corning 354235) followed by Trypsin-EDTA and Collagenase I treatment, expanded in 15-cm cell culture dishes, and frozen for further use. Progenitor cells were thawed and grown to100% confluency and then cultured in DMEM + 10% FBS supplemented with Isobutyl-3-methylxanthine (0.5 mM), Insulin **(5 µg/mL OR 8.6 nM)** and Dexamethasone (0.25 µM) (MDI) for 72 hrs. Subsequently, half of the medium was replaced with DMEM + 10% FBS every other day for 3 days. Experiments were performed at day 10 of differentiation unless otherwise noted. For immunofluorescence staining, cells were grown on glass coverslips in 24-well plate. For qPCR experiments, cells were grown in 24-well plates. For western blot analysis, cells were grown in 12-well plates.

### RNA extraction for bulk RNA-sequencing and qPCR

Cells in culture wells were washed with PBS before harvesting with TriPure TRIzol reagent (Cat# 11 667 165 001, Roche). The cell-TRIzol mixtures were transferred to collection tubes and homogenized with Tissuelyser II (Qiagen). Chloroform was added in a 1:5 ratio by volume and phase separation was performed. The RNA-containing layer was mixed with an equal volume of 100% isopropanol and incubated overnight at −20 °C for precipitation. RNA was pelleted and washed with 80% ethanol and resuspended in nuclease-free water. RNA concentration and purity were determined using a NanoDrop 2000 (Thermo Scientific). RNA for sequencing were sent to University of Massachusetts Medical School Molecular Biology Core Lab for fragment analysis.

### Bulk RNA-sequencing

Library preparation was performed using TruSeq Stranded mRNA Low-Throughput Sample Prep Kit (Cat# 20020594, Illumina) according to manufacturer’s instruction. The libraries were sequenced on the NextSeq 500 system (Illumina) using the NextSeq® 500/550 High Output Kits v2 (75 cycles; single-end sequencing; Cat# FC-404-2005, Illumina). The FASTQ files were processed using the DolphinNext pipeline [46] on the Massachusetts Green High Performance Computer Cluster (GHPCC). DolphinNext was configured to use RSEM for read mapping and transcript quantification [47]. Differentially expressed genes were identified using DESeq2 [48]. Pathway analysis was performed using ToppGene (Chen J, Bardes EE, Aronow BJ, Jegga AG 2009. ToppGene Suite for gene list enrichment analysis and candidate gene prioritization. Nucleic Acids Research doi: 10.1093/nar/gkp427 (PubMed)).

### RT-qPCR analysis

Total RNA was purified using Direct-zol RNA Miniprep Kit (Zymo Research) following the instructions of the manufacturer or using the TriPure Isolation Reagent. Reverse transcriptions were performed for 1 µg RNA using iScript cDNA Synthesis Kit (Bio-Rad 170-8891) according to the manufacturer’s protocol. cDNA was loaded in duplicate, and qPCRs were performed with iQ SYBR Green Supermix (Bio-Rad 170-8882) using CFX96 Touch Deep Well Real-Time PCR Detection System (Bio-Rad). The qPCR primers used in this study are listed in the following table.

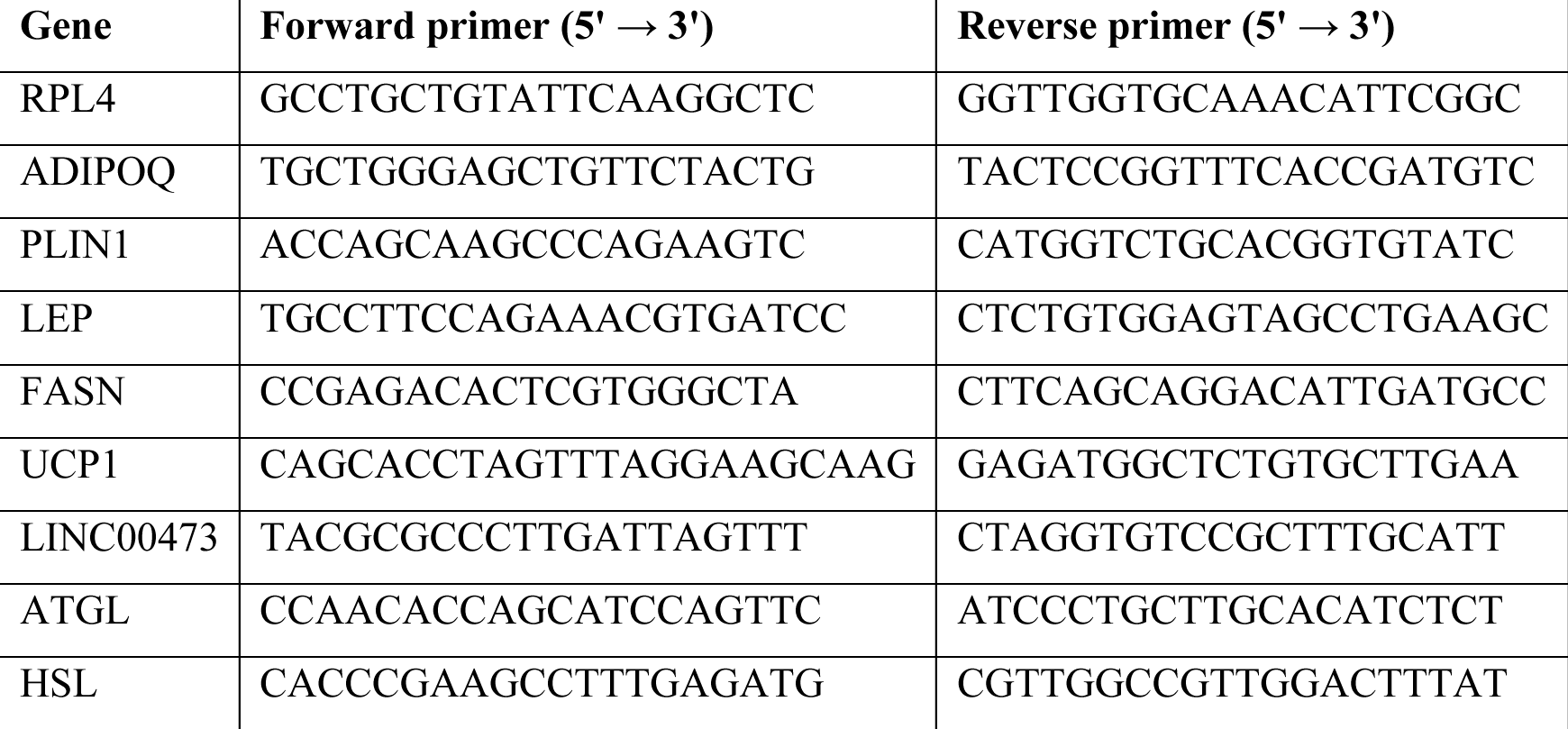

### RNAscope

*UCP1* mRNA was visualized using the RNAscope Multiplex Fluorescent Detection Kit v2 (Cat# 323100, ACD Bio) using target probes to *UCP1*(Cat#475821, ACD Bio) and labeled with Opal Fluorophore Reagents (Cat Akoya Biosciences) according to the manufacturer’s protocol. Signals were visualized and captured using a Zeiss Axiovert 200M Fluorescence microscope. ImageJ (FIJI) was used for image processing and normalization. All images analyzed were filtered to remove background using identical parameters, and values were expressed as a function of the number of cells in the field, assessed by Hoescht staining.

### Immunofluorescence staining

Cells were washed with PBS once, fixed with 4% paraformaldehyde in PBS for 15 min at room temperature and washed with PBS three times. Cells were permeabilized and blocked in PBS containing 1% BSA and 0.3% Triton X-100 for 30 min at room temperature and incubated with primary antibody in permeabilization and blocking buffer overnight at 4°C. Cells were washed with PBS three times. Secondary antibodies conjugated to Alexa Fluor-568 or Alexa Fluor-488 were used at 1:1000 dilution for 1 hour at room temperature. For nuclear staining Hoechst 33342 was used in permeabilization and blocking buffer for 5 min. Coverslips were mounted using Prolong Gold Antifade Mountant and imaged using Zeiss LSM 800 or Zeiss Axiovert 200M Fluorescence microscope. Images were analyzed using ImageJ (FIJI).

### Immunoblotting

Cultured adipocytes were washed with PBS twice at room temperature, and 0.12 ml of boiling 2% SDS solution supplemented with 1X Halt protease and phosphatase inhibitor cocktail was added to each well of a 12-well plate. Cell lysates were scrapped into 2-ml Eppendorf tubes and sonicated for 30 sec with 15 sec on/off cycles at 60% amplitude on ice. Protein concentrations of cell lysates were determined using BCA assay. 40 µg of whole cell lysates were combined with Laemmli buffer, heated at 95°C for 10 min and separated on SDS-PAGE gels. Proteins were transferred to PVDF membranes. Membranes were blocked in TBST containing 5% (w/v) skim milk for 1 hour at room temperature and incubated with primary antibodies in TBST containing 5% BSA at 4°C overnight. Primary antibodies were detected with HRP-conjugated secondary antibodies diluted in TBST containing 5% BSA at room temperature for1 hour. HRP activity was visualized after incubation of SuperSignal West Pico PLUS substrate and imaging on Bio-Rad ChemiDoc imaging system. The following antibodies and reagents were used: anti-UCP1 (Abcam ab10983, 1:200 dilution) anti-PLIN1 (Cell Signaling Technology 9349S, 1:1000 dilution), anti-Phospho-PLIN1-Serine 522 (Vala Sciences 4856, 1:4000 dilution), anti-HSL (Cell Signaling Technology 4107S, 1:1000 dilution), anti-ATGL (Cell Signaling Technology 2138S, 1:250 dilution), goat anti-rabbit HRP conjugated secondary antibody (Invitrogen 31480, 1:2500 dilution for HSL and ATGL blots, 1:10000 dilution for other immunoblots), goat anti-mouse HRP conjugated secondary antibody (Invitrogen 31430, 1:10000).

### CRISPR-Cas9 knockouts

Neon Transfection System 10 µL Kits (ThermoFisher MPK1025) were used for ribonucleoprotein (RNP) nucleofection according to the manufacturer’s protocol. Briefly, RNP complexes consisting of 4 µM sgRNA (IDT or Synthego) and 3 µM SpyCas9 protein (PNA Bio) was prepared in Buffer R provided by the Neon Transfection System Kit to a volume of 6 µl. Progenitor cells were trypsinzed, washed and resuspended in Buffer R to a density of 2×10^5^ cells per 6 µl. Equal volume of RNP mix and cell suspension were mixed using 10 µl Neon tip and electroporated using the optimized parameters (voltage: 1350V, width of pulse: 30 ms, number of pulse: 1). After electroporation, the cells were plated immediately in 12-WP containing 1 ml of complete media, grown to confluence, and differentiated to adipocytes according to the methods mentioned in <u>cell culture from human adipose tissue explants</u> for downstream experiments. The sequences of sgRNAs used in this study are listed in the following table.

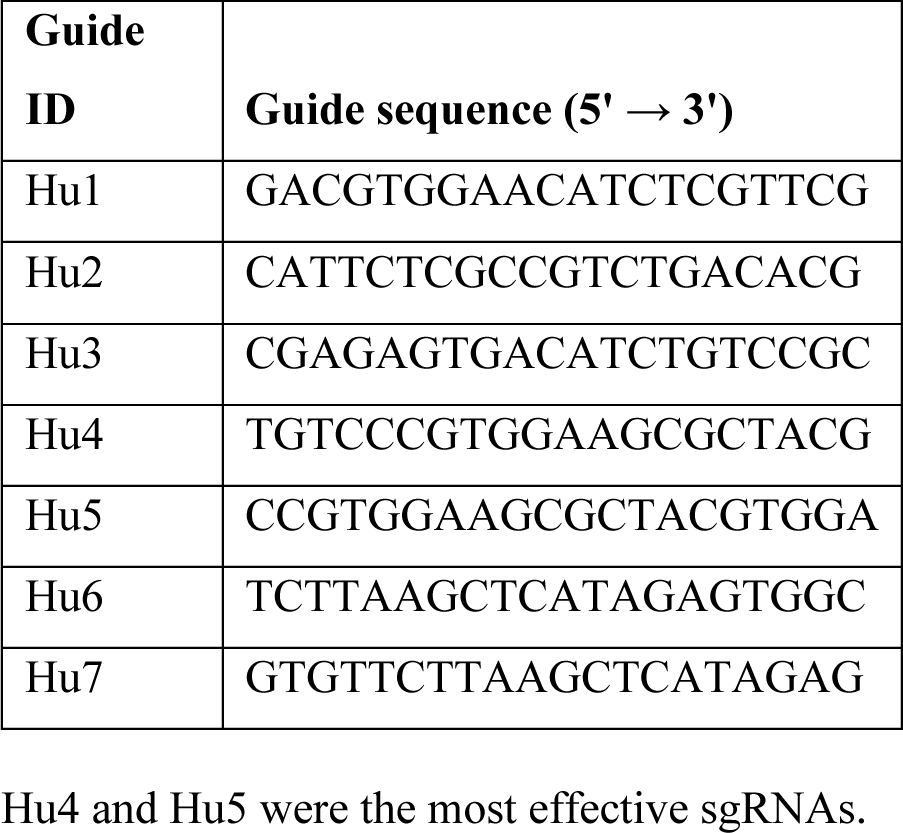

### Indel analysis by TIDE

The electroporated progenitors were subjected to adipogenic differentiation for 8 days and the genomic DNA was isolated using QuickExtract DNA Extraction Solution (Lucigen QE09050). PCR reactions were conducted with 100 ng of genomic DNA and following primer pairs spanning the sgRNA target site in KAPA HiFi HotStart ReadyMix (Roche KK2602) according to the manufacturer’s instructions. The PCR products were purified by QIAquick PCR Purification Kit (28106) and submitted to Genewiz for Sanger Sequencing. The control and sample sequence data files were analyzed using TIDE webtool (htEtps://tide.nki.nl/#about) to quantify indel frequencies. The PCR primers designed for sgRNAs targeting ATGL are listed in the following table.

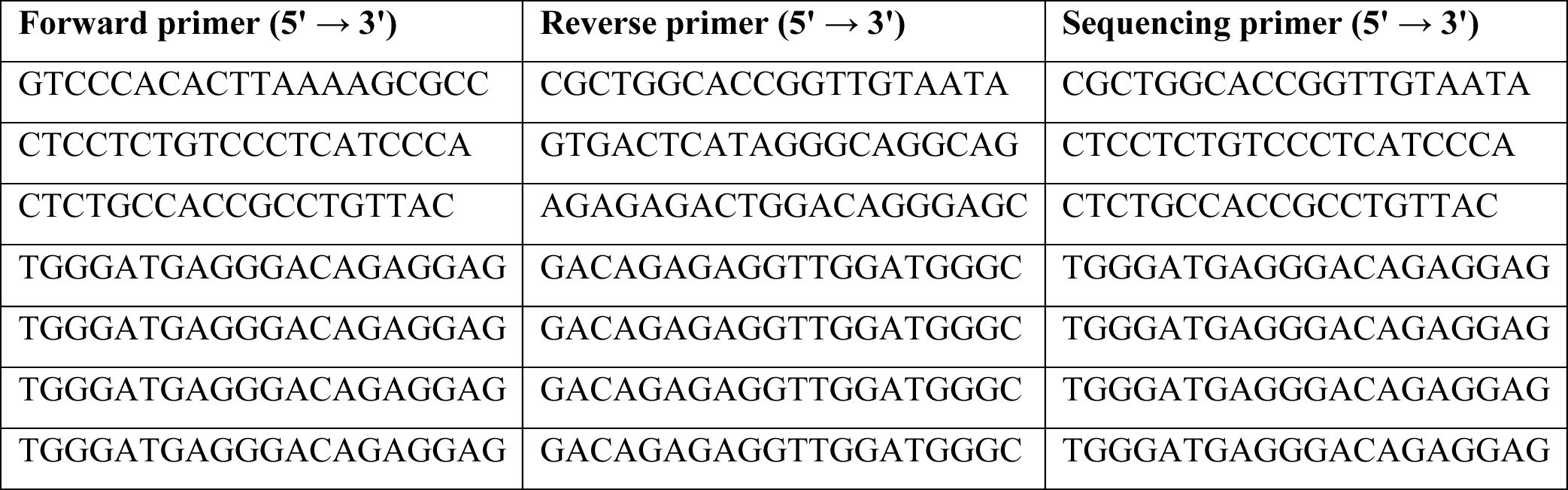

### Glycerol assay

Cell culture media were collected at specific time points and glycerol in media was measured using free glycerol reagent (Sigma-Aldrich F6428) according to the manufacturer’s protocol.

## ACKNOWLEDGEMENTS

This study was supported by NIH grants DK089101 and DK123028 to SC.

## DATA AVAILABILITY STATEMENT

All sequencing data will be deposited in a publicly accessible repository upon acceptance. All reagents generated for this project will be made available on request.

